# Protist.guru: a comparative transcriptomics database for the protist kingdom

**DOI:** 10.1101/2021.08.04.455173

**Authors:** Erielle Marie Fajardo Villanueva, Peng Ken Lim, Jolyn Jia Jia Lim, Shan Chun Lim, Pei Yi Lau, Kenny Ting Sween Koh, Emmanuel Tan, Ryanjit Singh Kairon, Wei An See, Jian Xiang Liao, Ker Min Hee, Varsheni Vijay, Ishani Maitra, Chong Jun Boon, Kevin Fo, Yee Tat Wang, Ryan Jaya, Li Anne Hew, Yong Yee Lim, Wei Quan Lee, Zhi Qi Lee, Herman Foo, Adriana Lopes dos Santos, Marek Mutwil

**Author notes:** Corresponding author: Marek Mutwil, School of Biological Sciences, Nanyang Technological University, 60 Nanyang Drive, 637551, Singapore, Singapore.

## Abstract

**Summary:** During the last few decades, the study of microbial ecology has been enabled by molecular and genomic data. DNA sequencing has revealed the surprising extent of microbial diversity and how microbial processes run global ecosystems. However, significant gaps in our understanding of the microbial world remain, and one example is that microbial eukaryotes, or protists, are still neglected. To address this gap, we used gene expression data from 15 distinct protist species to create protist.guru: an online database equipped with tools for identifying functional co-expression networks, gene families, and enriched gene clusters. Here, we show how our database can be used to reveal genes involved in essential pathways, such as the synthesis of secondary carotenoids in *Haematococcus lacustris*. We expect protist.guru to serve as a valuable resource for protistologists, as well as a catalyst for discoveries and new insights into the biological processes of microbial eukaryotes.

**Availability:** The database and co-expression networks are freely available from http://protist.guru/. The expression matrices and sample annotations are found in the supplementary data.

## Introduction

Microbial eukaryotes, or protists, are a phylogenetically broad group of single-celled organisms. The term “protist” was employed in the 19th century by the artist and biologist Ernest Haeckel (Haeckel, 1866) to all eukaryotes that were not plants, animals, or fungi. Today, we know that although most of the described species of eukaryotes belong to the multicellular groups of animals (Metazoa), plants, and fungi, these lineages only represent a very small proportion of the eukaryotic diversity (Ibarbalz *et al.*, 2019; Mahé *et al.*, 2017). Together with bacteria and fungi, protists form the engine of every biogeochemical cycle central to all ecosystems on earth. Through their microbial processes, they drive the cycling of nutrients and the energy flow between all planet spheres (e.g., biosphere and atmosphere).

Despite their impact on human health and on a planetary scale, the understanding of gene function in protists has lagged behind other microbial taxa. The reasons for this are numerous and range from technical challenges to a lack of readily cultivable strains. In addition, protists have much larger genomes and more complicated gene expression patterns when compared to bacteria. These features have led to minimal knowledge about the gene number, identity, and function within several protistan lineages (del Campo *et al.*, 2014). Since gene sequence alone can only result in a partial prediction and is not useful for genes that do not show sequence similarity to characterized genes (Rhee and Mutwil, 2014), methods based on gene expression have increasingly been used to predict gene function. Co-expression analysis finds functionally related genes by identifying genes that exhibit similar expression profiles across different growth conditions and genotypes (Mutwil *et al.*, 2011).

To improve our understanding of protists’ gene functions and expressions, the protist.guru database was constructed based on 1,651 transcriptomes. Protist.guru is equipped with a plethora of tools, empowering users to analyze gene expression profiles and co-expression networks across 15 protist species and allowing for the study of novel genes essential for biological processes in protists.

## Implementation

Our database has multiple tools to allow for different analyses of the genomic and transcriptomic data. The data can be viewed with ease through pages such as species (www.protist.guru/species), genes (www.protist.guru/sequence/view/48072), gene families (www.protist.guru/family/view/186501), co-expression clusters (www.protist.guru/cluster/view/600), neighborhoods (www.protist.guru/network/graph/39535), family phylogenetic trees (www.protist.guru/tree/view/48179), Pfam domains (www.protist.guru/interpro/view/13405), and Gene Ontology terms (www.protist.guru/go/view/4133). Each page contains additional information relevant to the type of data being displayed. For instance, the gene pages contain information about cDNA and protein sequences, functional annotations, expression profiles, co-expression neighborhoods, and more. On the other hand, Gene Ontology pages show GO annotations, the genes in the 15 protists with the same GO term, and enriched co-expressed clusters for genes that have that particular GO term. The features page (www.protist.guru/features) lists a complete description of search functions and tools.

To exemplify how our tool can be used to uncover novel genes and conserved gene clusters in biosynthetic pathways, we analyzed the secondary carotenoid biosynthesis pathway in *Haematococcus lacustris* via co-expression analysis. *Haematococcus lacustris* is a unicellular freshwater microalga that is a rich source of astaxanthin, a highly valued red xanthophyll known for its potent antioxidant activity (Han *et al.*, 2019).

Since phytoene is the first carotenoid precursor for astaxanthin biosynthesis, we queried our database with phytoene desaturase, one of the first two fundamental enzymes that catalyzes the conversion of C40 phytoene to ζ-carotene, an essential precursor for beta carotene, and hence astaxanthin. Upon entering the key intermediary enzyme, phytoene desaturase (lcl|BLLF01000057.1_cds_GFH06801.1_1278) for astaxanthin production in *Haematococcus lacustris* into our database, we arrive at its gene page (https://protists.sbs.ntu.edu.sg/sequence/view/152450). Stress inducing conditions have been shown to increase the yield of astaxanthin in *Haematococcus lacustris* cells, by inducing a preferential morphological transformation of vegetative, green, motile cells to mature, non-motile cysts filled with red astaxanthin (Minhas *et al.*, 2016). An overview of the gene expression of phytoene desaturase in different strains of *Haematococcus lacustris*, developmental stages and physicochemical conditions, such as high-light illumination across different time periods and in different acidic cultures, can be found on the gene page.

The phytoene desaturase gene (lcl|BLLF01000057.1_cds_GFH06801.1_1278) is also found in a co-expression cluster (Cluster 8: https://protists.sbs.ntu.edu.sg/cluster/view/2936), which represents groups of functionally related genes. The cluster page can be navigated from the gene page and displays a cluster expression profile that takes into account the TPM values of gene members across different conditions (Fig 1A). Information on significantly enriched Gene Ontology (GO) terms (corrected p-value <0.05), InterPro Domains, and gene families for gene members are also available. Based on the expression profile, we saw that the expression of the genes in the cluster showed the highest expression after 24 hours of high light exposure with a corresponding decrease in expression after 48 hours. This suggests that transient stress induction via high light exposure could be ideal to obtain the highest yield of astaxanthin. Thus, the expression profile analysis can reveal the conditions when a given gene and its associated pathway is highly expressed.

**Fig. 1.**
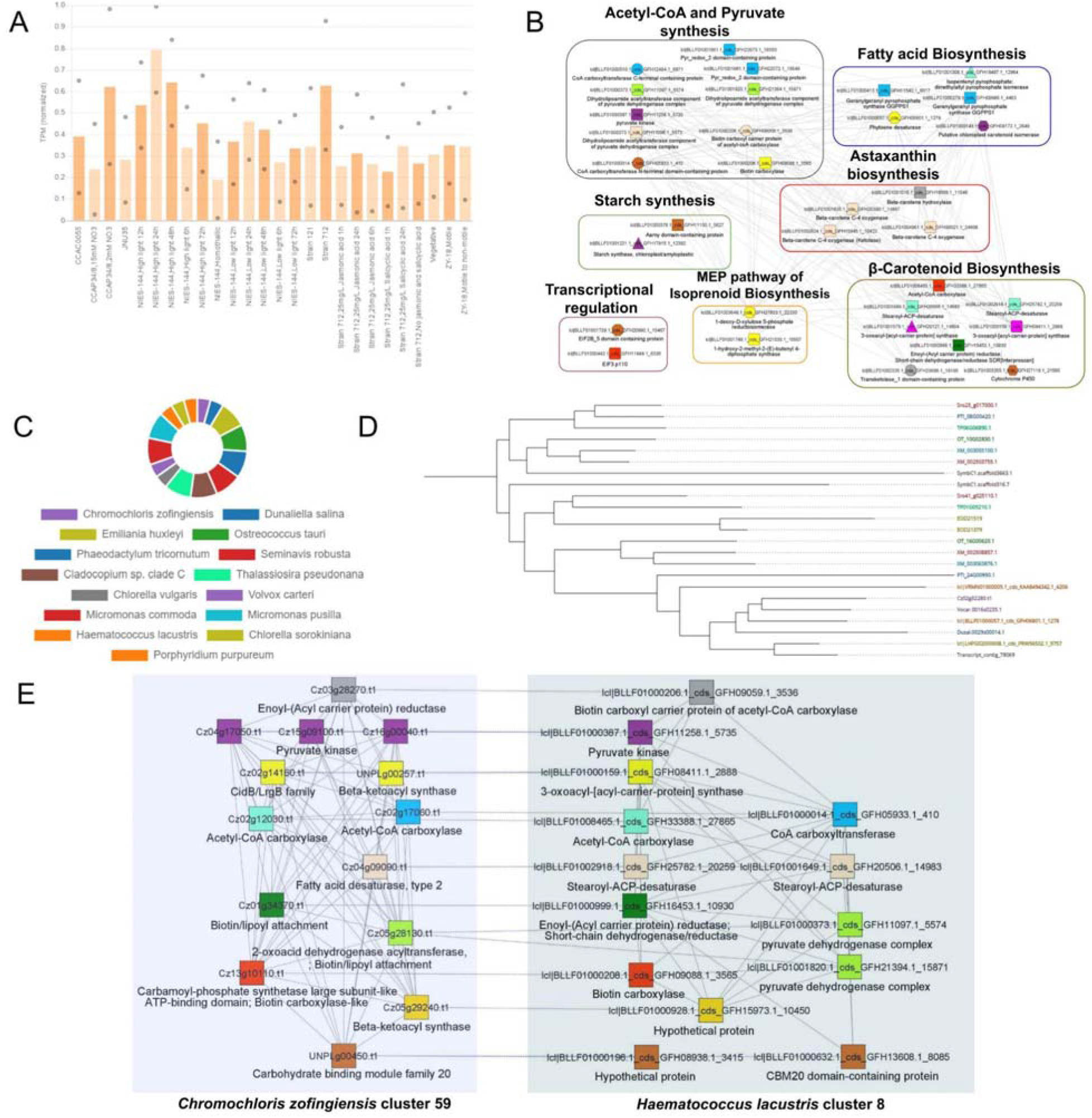
Outline of the tools available on Protist.guru exemplified with *Haematococcus lacustris*. A) Expression profile of cluster 8 with the average TPM values for all genes present in the cluster. Different strains and sample conditions are represented on the x-axis whereas gene expression values in Transcript Per Million (TPM) are represented on the y-axis. Dots and bars represent the maximum/minimum gene expression values and average gene expression values respectively. B) Co-expression network of genes identified in the same co-expression cluster as phytoene desaturase (lcl|BLLF01000057.1_cds_GFH06801.1_1278). Nodes represent genes and co-expressed genes are connected by edges in the network. Novel genes encoding for hypothetical proteins and proteins with generic pfam domains were removed from the network. Node colors indicate genes encoding for common Pfam domains and orthogroups. C) Proportion of sequences in OG_03_0001542 gene family across all 15 protists. A total of 23 sequences across 15 protists were identified. D) Phylogenetic tree for OG_03_0001542 gene family across 15 protists in the database. E) Comparative gene co-expression networks for similar clusters in *Haematococcus lacustris* and *Chromochloris zofingiensis.* Only genes that have a common homolog or Pfam domains are shown. For brevity, repetitions in gene descriptions were trimmed. Solid lines denote co-expressed genes whereas dashed lines denote homologous genes across the 2 clusters.

The co-expression cluster consists of 76 genes that are involved in various processes such as acetyl-CoA and pyruvate synthesis, methylerythritol phosphate pathway (MEP) of isoprenoid biosynthesis, fatty acid biosynthesis, β-carotenoid biosynthesis, astaxanthin biosynthesis, transcriptional regulation and starch synthesis (Fig 1B). The presence of these processes in the cluster implies functional relevance between the processes. Acetyl-coA and pyruvate are essential first precursors for carotenoid biosynthesis, which could be fed into mevalonate (MVA) or MEP pathway for isopentenyl pyrophosphate (IPP) and dimethylallyl pyrophosphate (DMAPP) synthesis. IPP and DMAPP are precursors for synthesis of terpenoids such as β-carotene, which is an intermediate for astaxanthin production. Stoichiometric coordination and interdependence between fatty acid biosynthesis and astaxanthin production pathways were also reportedly observed in *Haematococcus lacustris* with some fatty acid biosynthesis acyltransferases postulated to be involved in astaxanthin esterification (Chen *et al.*, 2015). Through co-expression analysis, the genes identified using our tool could provide greater insights into potential crosstalk between pathways that could affect astaxanthin biosynthesis in *Haematococcus lacustris*. This could be immensely useful for the metabolic engineering of *Haematococcus lacustris* cells to increase astaxanthin production.

Additionally, information on similar clusters from all species is available from the cluster page and can be compared using the “compare” button. Here, we present a similar cluster to *Haematococcus lacustris*’s cluster 8 in another astaxanthin producing microalga, *Chromochloris zofingiensis* with a Jaccard index of 0.094. Upon clicking “Compare” to compare the clusters, we arrive at a page showing the co-expression network comprising genes in the conserved clusters in the two organisms (Fig 1E). The genes conserved between these clusters are involved in acetyl-CoA, pyruvate, and fatty acid biosynthesis (Fig 1E). As demonstrated in this example, our tool allows for the easy identification of novel genes in different biological pathways and functional orthologs through comparing conserved neighbourhoods with genes in the same orthogroup and with similar co-expression.

## Conclusion

We developed protist.guru, a comparative transcriptomics database with multiple tools to analyze and visualize gene co-expression networks and expression profiles among 15 protists. Through this database, we hope to aid the investigation of gene function in protists and facilitate a deeper understanding of these microbial eukaryotes.

## SUPPLEMENTARY DATA

### Supplemental Figure

**Figure S1. Quality control and data selection for the 15 protists.** The number of processed reads (NPR, x-axis) versus the percentage of pseudoaligned reads (PPR, y-axis) are shown for all 15 species. Each dot represents one sample, where black and green indicate samples that failed and passed the quality control, respectively. The histograms to the right and on top of each plot indicate the distributions of the PPR and NPR values, respectively.

### Supplementary Methods

Tools and analyses used to construct protist.guru

### Supplemental tables

Due to the large size of the tables (∼300Mb), they can be downloaded from https://doi.org/10.6084/m9.figshare.15097419.v1

**Table S1**. Genome versions used to build the database

**Table S2.** Pseudoaligning statistics of the analyzed species

**Table S3**. Expression profile of CHLSO genes across RNA-Seq experiments

**Table S4.** Expression profile of CHLVU genes across RNA-Seq experiments

**Table S5.** Expression profile of CHRZO genes across RNA-Seq experiments

**Table S6.** Expression profile of CLADO genes across RNA-Seq experiments

**Table S7.** Expression profile of DUNAL genes across RNA-Seq experiments

**Table S8.** Expression profile of EMIHU genes across RNA-Seq experiments

**Table S9.** Expression profile of HAELA genes across RNA-Seq experiments

**Table S10.** Expression profile of MICCO genes across RNA-Seq experiments

**Table S11.** Expression profile of MICPU genes across RNA-Seq experiments

**Table S12.** Expression profile of OSTTA genes across RNA-Seq experiments

**Table S13.** Expression profile of PHATR genes across RNA-Seq experiments

**Table S14.** Expression profile of PORPU genes across RNA-Seq experiments

**Table S15.** Expression profile of SEMRO genes across RNA-Seq experiments

**Table S16.** Expression profile of THAPS genes across RNA-Seq experiments

**Table S17.** Expression profile of VOLCA genes across RNA-Seq experiments

