## Supplementary figures and images for "Protist.guru: a comparative transcriptomics database for the protist kingdom"

### Figure S1

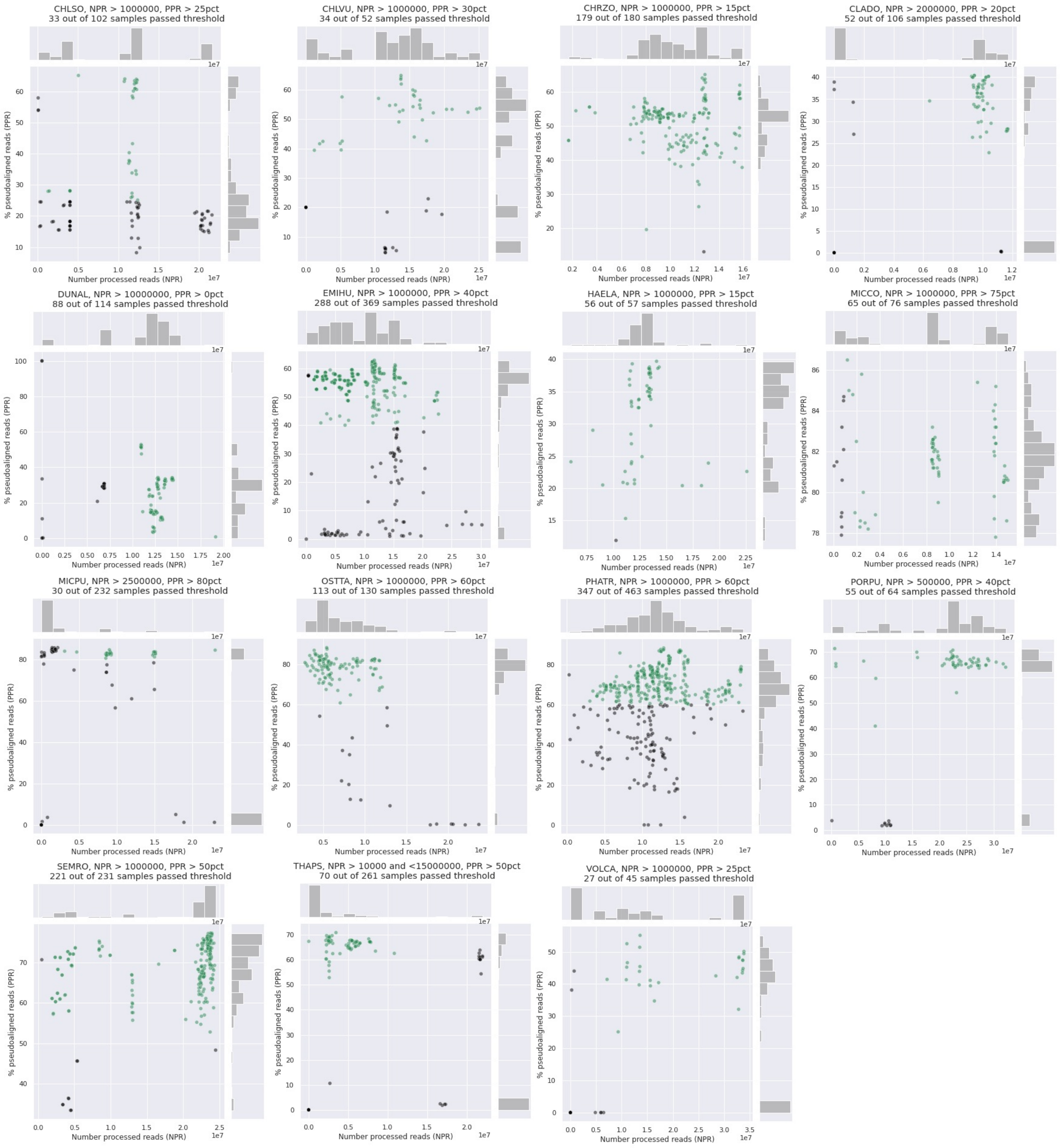
